# Repeated evolution of convergent iridescence in closely-related species of *Morpho* butterflies living in sympatry

**DOI:** 10.64898/2026.02.06.704458

**Authors:** Joséphine Ledamoisel, Vincent Debat, Violaine Llaurens

## Abstract

The evolution of visual traits in closely-related species living in sympatry is highly influenced by their ecological interactions: while sexual selection tends to promote the divergence of visual cues involved in mate choice, natural selection via predation may promote the convergence of dissuasive signals between prey species, especially in unpalatable or evasive prey. Here, we investigate the impact of sympatry on the evolution of the blue structural colouration in the wings of two closely-related *Morpho* butterfly species across several localities throughout Central and South America. Dorsal iridescence might affect mate choice and species recognition, which should promote its local divergence among species. However, the bright flashes and dynamic colour patterns produced by iridescence during flight might also increase survival by confusing predators and favouring escape. Such an effect might in turn lead to convergence in wing iridescence between evasive species occurring in sympatry, a phenomenon dubbed evasive mimicry. To test the effect of these putative antagonistic selective forces on visual cues evolution, we quantified the variation of the structural blue colour displayed at 13 different combinations of illumination/observation angles, on the wings of two closely-related *Morpho* species. We contrasted 10 sympatric and 11 allopatric locations and specifically compared the phenotypic distances between individuals from different species. Phenotypic distances between heterospecific pairs of individuals were significantly smaller in sympatry, consistent with the hypothesis of a local convergence of iridescence due to evasive mimicry. Interestingly, sexual dimorphism was found between males and females, suggesting that the trade-off between natural and sexual selection on the evolution of iridescence might differ between sexes. Our results suggest that local predation pressures may promote repeated evolutionary convergence of structural colouration between evasive prey species living in sympatry.

## Introduction

How much do traits exhibited by closely-related species evolve independently when they occur in sympatry? Closely-related species share a common phylogenetic history and are thus more likely to present similar suites of traits (Blomberg et al., 2003) favouring their co-existence within the same habitat (Burns & Strauss, 2011). However, niche overlap caused by their phenotypic proximity might trigger acute competition for resources or reproduction, promoting the divergent evolution of traits involved in foraging and mate recognition in sympatry (Pfennig & Pfennig, 2009). In particular, in animals where visual cues are involved in mate recognition, divergent evolution of visual traits among closely-related species is frequently observed in sympatry (Allen et al., 2014; Stanger-Hall & Lloyd, 2015). However, the evolution of visual traits in closely-related prey species living in sympatry might also depend on the perception and reaction of shared predators (Anderson & de Jager, 2020; Nosil & Crespi, 2006). Predation pressures can indeed promote either directional or diversifying selection acting on visual traits displayed in prey (Nokelainen et al., 2024). Shared predation pressures may for instance promote the evolution of mimicry among unprofitable prey species living in sympatry, making the evolution of visual traits in these different prey species non-independent from one another. Comparing the evolution of visual traits in closely-related species in sympatric *vs*. allopatric areas thus allows to investigate the relative contributions of shared selective pressures and species interactions on the evolution of traits in sympatry.

The non-independent evolution of colouration between closely-related species living in sympatry is documented in a large number of animal taxa. The evolution of colouration has been shown to be strongly influenced by selection exerted by local abiotic factors such as temperature (Clusella Trullas et al., 2007), humidity (Delhey, 2019) or UV-radiation (Bastide et al., 2014), resulting in similar colouration among sympatric species because of these shared selective pressures. Phenotypic evolution of closely related species may also be deeply impacted in sympatry by natural selection arising from ecological interactions both (1) among those closely-related species and (2) between these and other species from other guilds. Indeed, colouration is often involved in competitive interactions, mate recognition and mate choice, as well as predator deterrence or avoidance (Protas & Patel, 2008), generating antagonistic selection regimes acting on the evolution of colouration in closely-related species in sympatry. Reproductive interference (Gröning & Hochkirch, 2008) might promote divergence in colouration: for example, larger divergence in throat fan colourations has been documented among closely-related species of lizards in sympatric areas *vs*. allopatric areas (Lambert et al., 2013). In contrast, predation pressures shared by different prey species in sympatry have been shown to lead to convergent evolution of colouration in multiple taxa (Joron & Mallet, 1998). Unpalatable prey indeed frequently display conspicuous colouration visual signals (Caro & Ruxton, 2019) recognised and avoided by predators (Darst & Cummings, 2006; Ihalainen et al., 2007), therefore generating a positive density-dependent selection acting on visual trait evolution both within and among unpalatable prey species (Joron & Mallet, 1998). Such mimicry caused by predator learning has also been hypothesised to happen in palatable prey with high escape abilities (Loeffler-Henry et al., 2021; Pinheiro, 1996; Young, 1971). Behavioural experiments have indeed shown that birds that were repeatedly presented different types of coloured prey quickly stopped to attack evasive prey associated with specific conspicuous colouration; the association between evasiveness and colouration was even more rapidly learnt than when the same colouration was associated with unpalatability (Páez et al., 2021). This learning process could eventually lead to the evolution of similar conspicuous colourations associated with evasiveness (*i*.*e*. evasive mimicry, Linke et al., 2022; Ruxton et al., 2004), among individuals belonging to different species submitted to the same predator community. For example, beetles sharing similar coloured traits associated with evasive flight are less targeted by visual predators, suggesting that resembling to a *hard-to-catch* prey may improve survival (Guerra et al., 2024). Convergent evolution of colouration might thus be promoted in closely-related species living in sympatry when they exhibit high evasiveness.

Here, we investigate variations in iridescent colours within and among two closely-related species of *Morpho* butterflies living in sympatry, using evasive mimicry as a working hypothesis. We specifically focus on the sister-species *Morpho helenor* (Cramer, 1776) and *Morpho achilles* (Linnaeus, 1758). These neo-tropical species display large overlapping geographic distributions (Blandin & Purser, 2013), allowing to estimate the effect of ecological interactions on the evolution of their traits by comparing multiple sympatric and allopatric localities. They are characterised by their very similar conspicuous dorsal wing patterns composed of a bright blue band, generated by specific scale nanostructures (Giraldo et al., 2016), surrounded by a dark pigmentation enhancing contrast. The blue dorsal colouration varies depending on the angle of observation and illumination (Ledamoisel et al., 2025), thereby producing a dynamic iridescent visual phenotype during flapping flight. Male mate choice experiments using female wing models in the wild (Le Roy et al., 2021) and in controlled conditions (Ledamoisel et al., 2025) suggested that iridescent colouration may be involved in mate localisation and mate choice in these *Morpho* species. Interestingly, the ventral wing surface exhibits a dull brown pigmentary colouration contrasting with the bright dorsal colouration. The blue iridescence, coupled with their erratic flapping flight, thus generates a confusing visual signal: the alternation between the ventral matte and the dorsal bright blue sides of their wings during flight creates conspicuous flashes. Combined with their unpredictable flight trajectories, these flashes could make these butterflies very difficult to follow, locate and catch by bird predators (Murali, 2018; Young, 1971); Capture-Mark-Recapture experiments indeed showed that iridescent flashes were as efficient as cryptic colouration in predation avoidance (Vieira-Silva et al., 2024). In turn, these very conspicuous signals could make them highly memorable to predators, which would learn to refrain from attacking such hard-to-catch blue prey. In agreement with this hypothesis, observations in the field have shown that *Morpho* butterflies are often sight-rejected by wild insectivorous birds, as compared to alternative butterfly prey from the same community (Pinheiro & Campos, 2019). Overall, the evasive capacities of *Morpho* butterflies combined with their very conspicuous iridescent colouration could provide protection against predators, and promote the evolution of similar iridescent patterns in different species facing the same predator community (*i*.*e*. evasive mimicry). Because chemical defences have never been reported in those butterflies, and because their caterpillars feed on non-toxic Fabacea and bamboo hostplants (Blandin et al., 2014; Chazot et al., 2021), evasiveness rather than toxicity has been assumed to promote colour pattern similarity between sympatric species of *Morpho*. A previous study investigating variations in the width of the dorsal blue band in *M. helenor* and *M. achilles* indeed found that these species share more similar wing *patterns* when they live in sympatry compared to allopatry (Llaurens et al., 2021), consistent with the hypothesis of local convergence of wing pattern expected via evasive mimicry. Studying how much blue iridescence actually converges between *Morpho* species within locality thus allows to investigate the respective effect of visual interactions with predators (favouring convergence) and heterospecific mates (promoting divergence) in the evolution of iridescence pattern among closely-related species.

We used Museum specimens to quantify iridescence variation within and between the closely-related species *M. helenor* and *M. achilles* in both allopatric and sympatric ranges. We computed the phenotypic distances based on iridescence between heterospecific pairs of individuals across localities and tested whether these distances were lower among pairs sampled in the same locality compared to pairs collected in different localities. We then used visual modelling to test whether birds are able to perceive the differences in iridescence among species within localities. Altogether, our study aims at testing whether geographic variation in iridescent blue displayed in closely-related *Morpho* species is consistent with the pattern expected if evasive mimicry shapes the evolution of colouration in sympatry.

## Materials and Methods

### Sampling zones and Museum specimens

We sampled *Morpho* specimens stored in the Muséum National d’Histoire Naturelle (MNHN) in Paris (France). *Morpho* butterflies are distributed across Central and South America, from Mexico to North Argentina (Blandin & Purser, 2013). *Morpho helenor* contains 40 described subspecies with substantial wing colour pattern differences, and has the largest distribution out of the two species studied here, with geographic overlap with *M. achilles* throughout the Amazonian basin (Blandin 2007). *M. helenor* thus sometimes co-exist with *M. achilles* in sympatry in the same microhabitat (the forest understory) at the same period of the year, but is also found isolated in allopatric regions located in Mexico, Costa Rica, Argentina, Eastern Brazil, and also in some localities in Western Ecuador, Northern Peru and Venezuela (see Figure 1).

**Figure 1:**
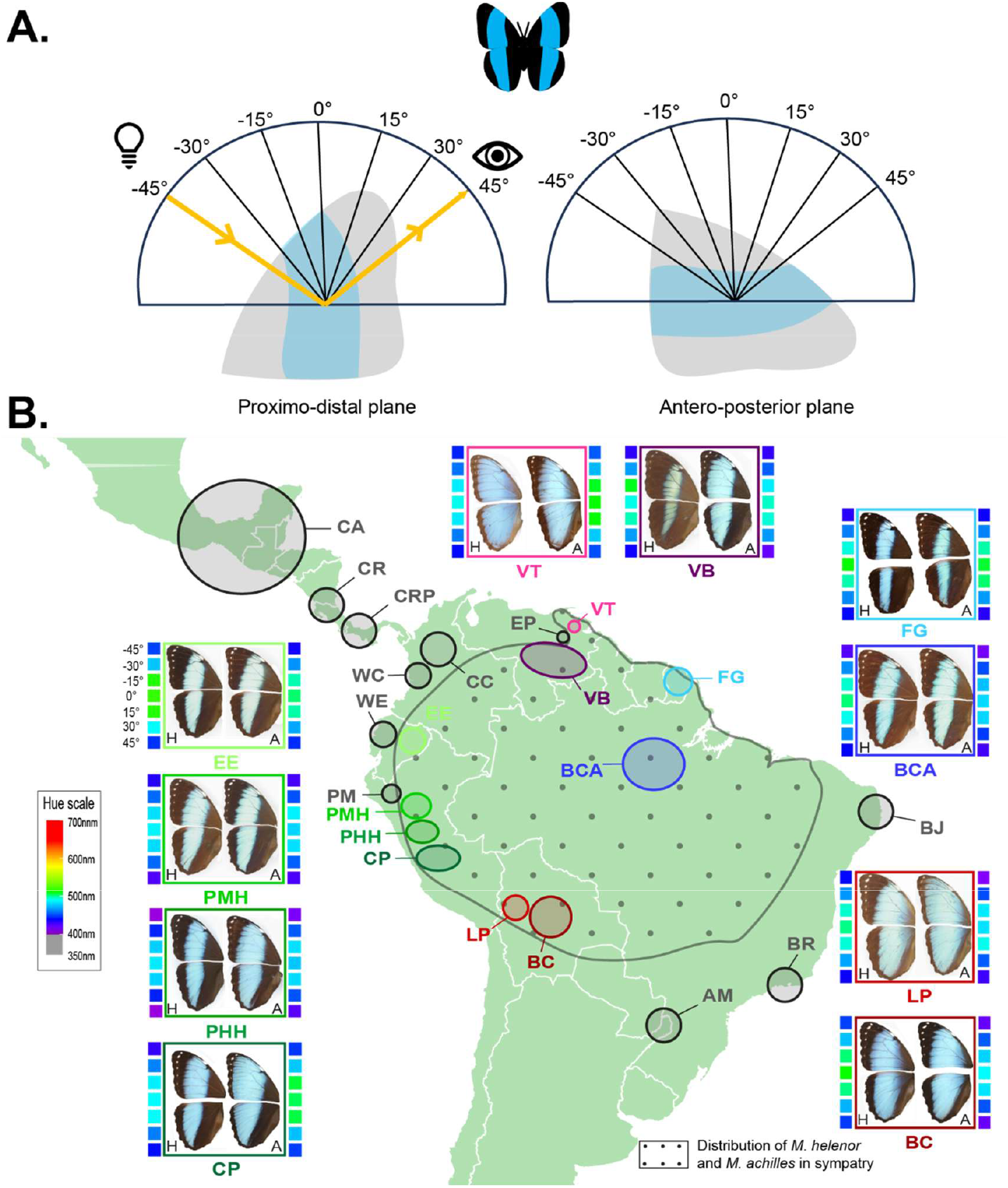
(A) Iridescence measurement protocol and (B) Geographic distribution of the *Morpho* samples studied, partitioned into 21 localities located across Central and South America, and their mean variation of colouration at different angles of illumination. (A) The specular reflectance of *Morpho* wings was measured when illuminating the wing along two planes (Proximo-distal and Antero-posterior planes) at 7 angles of illumination each, resulting in 13 reflectance spectra (the normal being shared by the two planes). Yellow arrowed lines show the position of illumination and observation fibers for a specular measurement of the reflectance of a wing in the Proximo-distal plane at a 45° angle as an example. (B) The black areas highlight the localities where *M. helenor* lives in allopatry from its closely-related species *M. achilles* (CA=Central America, CR=Costa Rica, CRP=Costa Rica Panama, CC=Central Columbia, WC=Western Columbia, WE=Western Ecuador, PM=Peru Maranon, AM=Argentina Missiones, BR=Brazil Rio de Janeiro, BJ=Brazil Joao, EP= Venezuela El Pao). The coloured areas indicate the localities where *M. helenor* lives in sympatry with *M. achilles*, either west (shades of green: EE=Eastern Ecuador, PMH=Peru Middle Huallaga, PHH=Peru High Huallaga, CP=Central Peru), south (shades of red: LP=Bolivia La Paz, BC=Bolivia Cochabamba), east (shades of blue: BCA=Brazil Central Amazon, FG=French Guiana) or north (shades of pink: VB=Venezuela Bolivar, VT=Venezuela Tucupita) of the Amazon basin. The pictures show the wing morphology of male *M. helenor* (H, on the left) and *M. achilles* (A, on the right) depending on the locality. Their respective hue variation measured along the Proximo-distal plan of their wings are pictured next to each subspecies: from top to bottom, each square represents the mean hue measured when illuminating the wings every 15° from −45° to 45°. Note that the entire hue variation at every angle of illumination measured on both wing planes, for each subspecies, sex, in every locality (allopatry or sympatry), can be found in Figure S1. This Hue representation follows an RGB scale: it is only illustrative and do not simulate the actual vision of neither butterflies nor predators.

Sampling zones were defined referring to *M. helenor* subspecies (Le Moult and Réal 1962, Blandin 2007). We used specimens collected in 21 localities across Central and South America, covering 18 *M. helenor* subspecies: subspecies distributed across large geographical zones (such as *M. helenor helenor, M. helenor theodorus* or *M. helenor coelestis*) were divided into different localities. Among those 21 localities, 10 are defined as *sympatric*. The remaining 11 localities are defined as *allopatric* for *M. helenor* because *M. achilles* does not occur in those regions. Using the same localities as in Llaurens et al. (2021) allowed us to test whether the convergence of the colour pattern (blue band width) found among sympatric species in this previous study extends to iridescence itself. We quantified the wing colour variations of *n* = 351 specimens in total, including *n* = 249 *M. helenor* and *n* = 102 *M. achilles* butterflies and used both males (*n* = 201) and females (*n* = 150). Females are much rarer than males in collections because of behavioural differences between sexes: *Morpho* males are more often observed flying along forest trails as compared to females and are thus more often caught by humans (see Table S1 for the detailed sampling plan).

### Iridescence measurement protocol

Iridescence is defined as the variation of colouration of a surface depending on the angle of illumination and /or the angle of observation. The reflectance of *Morpho* wings was thus measured at different angles of illumination and observation using a specular reflection set-up (*i*.*e*. by positioning the lightening and receiving probes symmetrically relative to the normal to the wing surface), following the protocol described in Ledamoisel *et* al. (2025). Briefly, wings were positioned on a flat surface, following a template to ensure that their orientation remained the same across all measurements. Iridescence was measured in the wing area delimited by the M3 and Cu1 veins using a spectrophotometer (AvaSpec-ULS2048CL-EVO-RS 200-1100nm, Avantes) coupled with a deuterium halogen light source (AvaLight-DH-S-BAL, Avantes), and two optical fibers (FCR-7UVIR200-2-1.5X100 and FC-UVIR200-2-1.5X100, Avantes). The angular position of the two fibers was precisely controlled using an angled fibre holder (AFH-15, Avantes). Reflectance was measured every 15° in the proximo-distal plan of the wings, the normal to the wing being set at 0°, resulting in 7 angular positions (−45°, −30°, −15°, 0°, 15°, 30°, 45°). The same protocol was repeated in the antero-posterior plan, resulting in a total of 13 reflectance spectra ranging from 300 to 700nm, measured at 13 angular positions per individual (the normal being common to the two plans, see Figure 1.A).

Although the repeatability of this protocol was validated in Ledamoisel et al. (2025), we repeated each measurement 3 times at every angle of illumination for 10 individuals. Repeatability was tested using the R package rptR 0.9.22 (Stoffel et al., 2017): no significant difference was detected between replicates for any of the tested angles, highlighting the robustness of the measurements.

### Spectral data analysis and statistics using colorimetric variables

#### Iridescence description

We used the R package pavo2 2.9.0 (Maia et al., 2019) on each of the 13 reflectance spectra measured per individual (*n* = 351) to compute the three colorimetric variables most commonly used to describe visual signals: Hue (*i*.*e*. the wavelength corresponding to the maximum reflectance, describing colour), Brightness (*i*.*e*. the mean reflectance over a spectrum, describing the amount of reflected light) and Chroma (*i*.*e*. the ratio of the reflectance range of each spectra divided by the brightness, describing colour saturation). To focus on the variation of blue iridescence, these colorimetric variables were extracted on the [350-700 nm] range from the spectra measured on the [300-700 nm] range. Because the normality and heteroscedasticity assumptions were rarely met for the distribution of values in these three parameters, we used permutation-based ANOVAs (vegan 2.6.4 package, Oksanen et al., 2001) to test the effect of sex, species, angle of measurement and the interactions between those variables on each wing plane separately and within each locality.

To investigate the variation of iridescence among individuals, we performed a Principal Component Analysis (PCA) based on Hue, Brightness and Chroma from the 13 reflectance spectra per individual. Each dot in the PCA space thus corresponds to the summary of reflectance spectra at different angles of an individual wing: the closer the points, the more similar the iridescence between individuals. The first 10 PCs (representing more than 98% of the explained variance) were retained and used in PERMANOVA to test for the effect of species, sex, and locality on iridescent patterns.

#### Testing iridescence convergence

To test whether individuals from either species were more similar within each sampling zone than expected by chance, we used the permutation-based approach previously described in Llaurens *et* al. (2021). The phenotypic resemblance between species was computed as the average Euclidean distances between individuals across species and localities, in the PCA space defined by the first 10 PCs computed from the Hue, Brightness and Chroma variations across angles. To test whether these phenotypic distances between individuals from different species are smaller when individuals are sampled within the same geographic areas as compared to randomly sampled individuals, a null distribution of phenotypic distance was built by randomly shuffling the location of the different individuals. For 10,000 permutations, the average phenotypic distance between interspecific pairs of *M. helenor* and *M. achilles* individuals was computed. A null distribution of phenotypic distances was thus generated assuming that geographic variation in colour pattern evolved independently in *M. helenor* and *M. achilles*. The significance of the phenotypic convergence among species in sympatry was assessed by comparing the observed average distance between interspecific pairs of individuals in sympatry to this null distribution: an observed distance was considered significantly smaller (*i*.*e*. greater phenotypic resemblance) when it fell below 95% of the randomised values.

We also used the PCA scores to estimate the levels of *M. helenor* intra-specific phenotypic variation when living in sympatry with *M. achilles* vs. allopatry. Under the hypothesis of evasive mimicry, the phenotypic variance within population is expected to be lower in sympatric than in allopatric populations of *M. helenor*, as a result of selection favouring resemblance among species. The phenotypic variance is expected to be similar in sympatric populations of the two species. Within population variance was computed as the trace of the covariance matrix of the PC scores within species within locality, for males and females separately.

### Spectral data analysis and statistics using “whole reflectance” dataset

As iridescence refers to any change in reflectance depending on illumination and observation angles, summing up iridescent signals with only 3 colorimetric variables from the [350-700 nm] range does not capture the whole variation in the visual signal, especially in the UV range. We thus also characterised the iridescence of each individual using the whole spectra obtained at all angular positions, by considering the reflectance at each wavelength as a different variable (400 wavelengths x 13 angular positions). We applied the same protocol as described above on this dataset: we performed a PCA, and retained the first 10 PCs to test for the effect of species, sex, and locality on iridescence using a PERMANOVA. The permutation-based approach to test for the convergence of iridescent signal between sympatric populations was also applied to the distances computed in this 10 PCs space depicting the full reflectance variation.

### Visual modelling

We tested whether the differences in iridescence detected between individuals from different species could be perceived by insectivorous birds using the R package pavo2 2.9.0 (Maia et al., 2019). Indeed, if convergent wing visual signals have been promoted by escape mimicry, predators should generalise the visual signals displayed by individuals from different species in sympatry, likely not perceiving subtle differences in iridescence between them. We specifically used the avian UV-sensitive vision model implemented in pavo (“avg.uv”) as a proxy for predator vision. Among each locality, the average chromatic (*dS*) and achromatic (*dL*) distances between species and sexes (respectively testing for hue and brightness discrimination) were calculated. When *dS* ≥ 1 and *dL* ≥ 1 Just Noticeable Difference (JND), we considered that the differences in blue could be perceived by an avian predator: this highly sensitive threshold is conservative regarding the discrimination capacities of predators.

## Results

### Variation in wing iridescence

By estimating the Hue, Brightness and Chroma from the reflectance spectra obtained at 13 different angles for each individual (Figure 1.A), we first compared the iridescent signal displayed in different populations of *M. helenor* and *M. achilles* (see Figure S1 showing the variation of Hue measured at different angles of illumination in every sympatric and allopatric locality).

The PCA performed on the Hue, Brightness and Chroma measured at every angles of illumination then allowed to summarise the combined effects of these colorimetric variables. According to factor loadings, Hue variables highly contribute to PC1 and 2 (see Figure 2): angular variation in Hue thus appears to be an important driver of the iridescent signal among *Morpho* individuals (Figure S2). The PERMANOVA based on 999 permutations performed on the first 10 PCs shows that iridescence is significantly different between species (*df* = 1, *F* = 27.430, *p-value* = 0.001), sexes (*df* = 1, *F* = 13.718, *p-value* = 0.001) and localities (*df* = 20, *F* = 9.462, *p-value* = 0.001). Additionally, the interactions between species and sex (*df* = 1, *F* = 10.965, *p-value* = 0.001), species and locality (*df* = 9, *F* = 1.944, *p-value* = 0.005) and sex and locality (*df* = 20, *F* = 1.718, *p-value* = 0.001) are also significant. These results highlight the great variation of wing iridescence across Central and South America (see Figure 3.A), but also between *Morpho* species. Note that the PCA based on the whole reflectance spectra dataset showed similar results (Figure S3).

**Figure 2:**
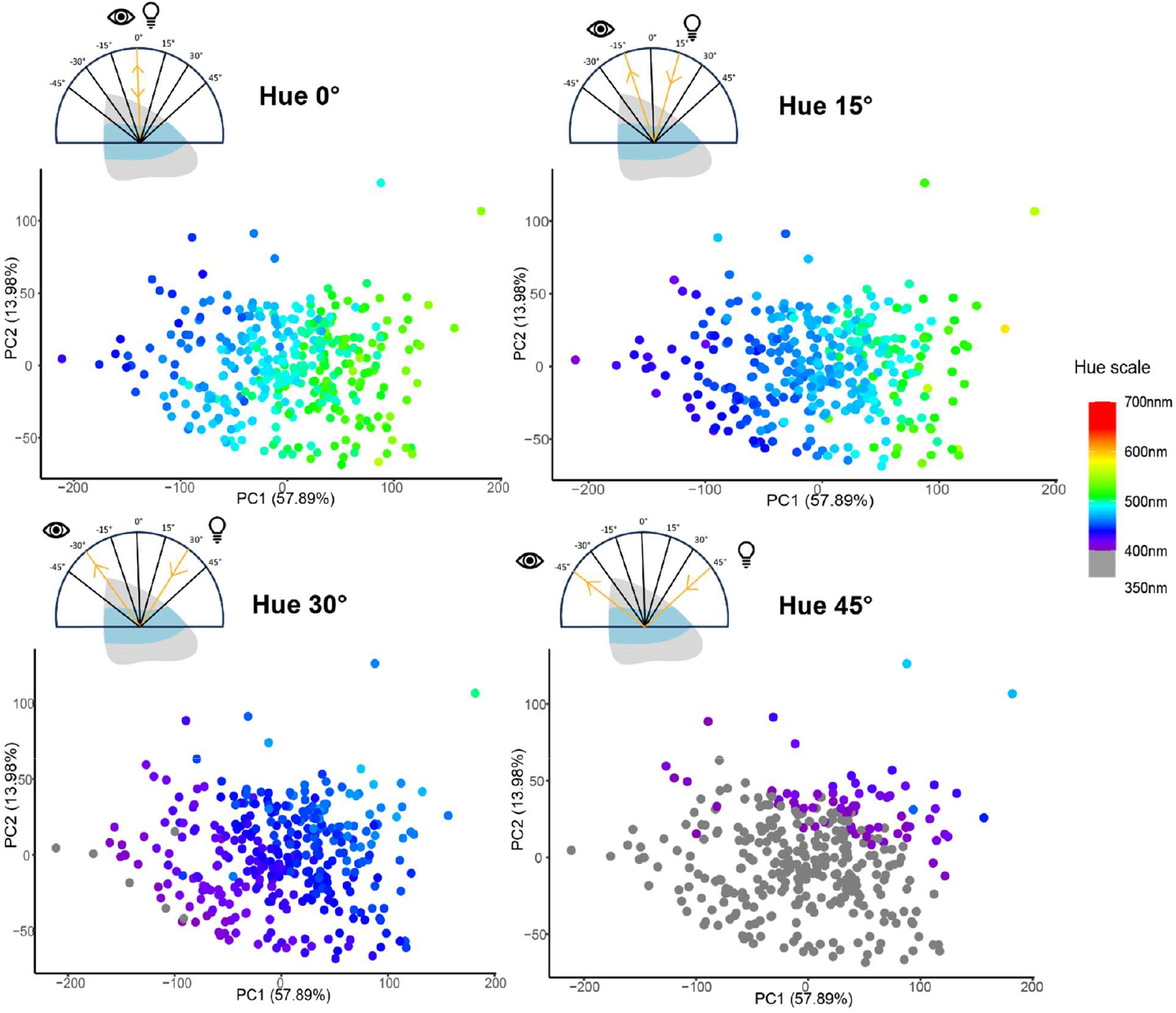
Principal component analysis based on the Hue, Brightness and Chroma variables measured at every angle of illumination on both wing planes of *Morpho* specimens. Each point on this graph represents the global signal of colorimetric variables depending on the angle of illumination (*i*.*e*. iridescence) of a given individual: the closer the points are together, the more similar the iridescence of two individuals is. Note that PC1 and PC2 account for more than 70% of the variance, and that both axes are correlated to Hue variables at different angles of illumination (see Figure S2). Here, we coloured each individual dot with their hue value at a 0° angle of illumination, and at 15°, 30° and 45° on their anterior side of their wings respectively.

**Figure 3:**
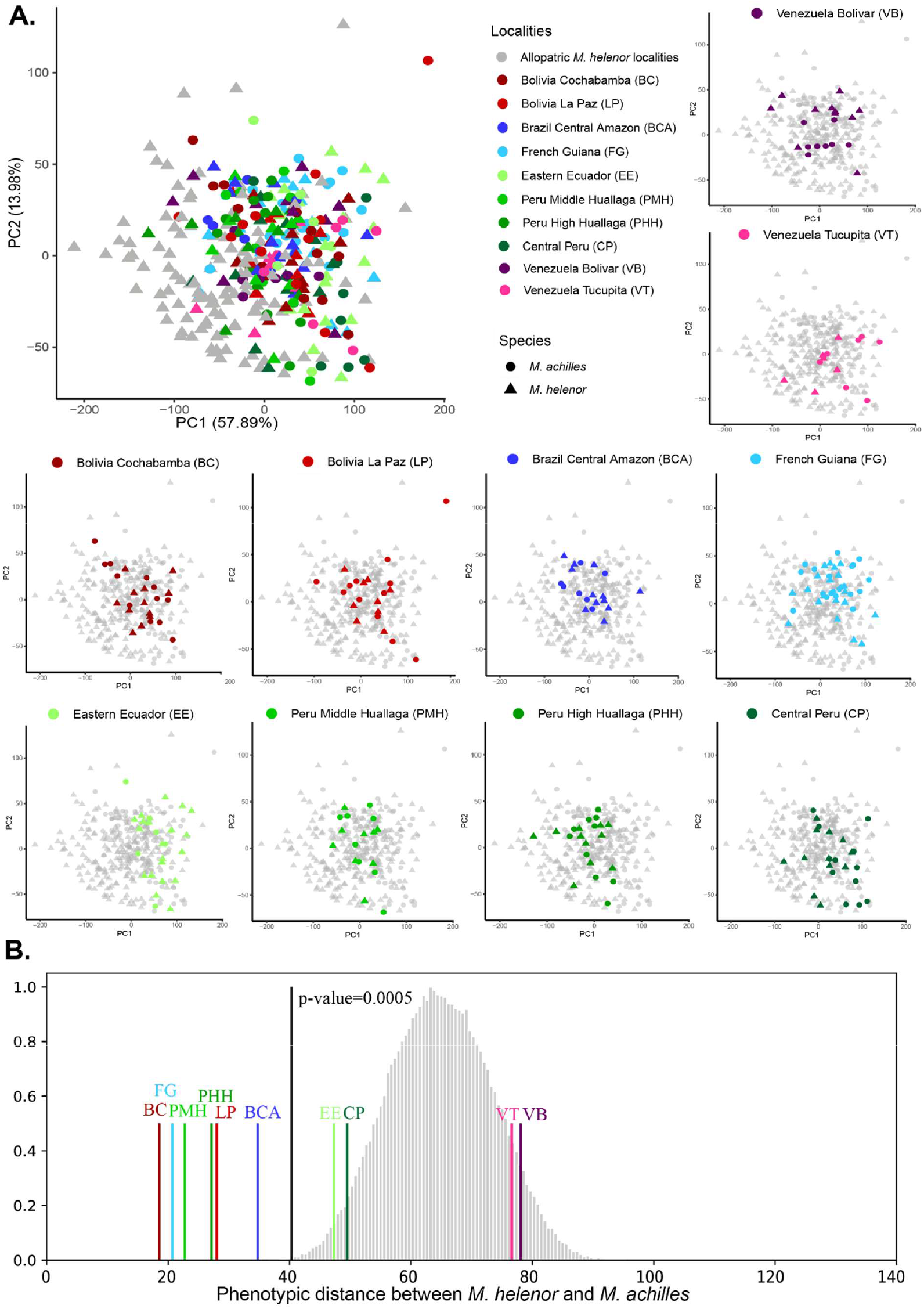
Phenotypic distances between *M. helenor* and *M. achilles* sampled within different geographic regions. (A) Principal component analysis based on the Hue, Brightness and Chroma variables measured at every angle of illumination on both wing planes of *Morpho* specimens. Each point on this graph represents the global signal of colorimetric variables depending on the angle of illumination (i.e. iridescence) of 1 individual: the closer the points are together, the more similar the iridescence of two individuals is. Grey points represent the iridescent signal of allopatric *M. helenor* found living in allopatry, while *Morpho* individuals sharing their habitat with closely-related species are highlighted in colours according to their locality shown on Figure 1. *M. achilles* individuals are represented with circles, *M. helenor* with triangles. Note that PC1 and PC2 account for more than 70% of the variance. (B) Phenotypic distances between *M. helenor* and *M. achilles* within different geographic regions (coloured bars) the predicted null phenotypic distribution between species obtained using 10,000 bootstraps, randomly reallocating the different sampling localities among individuals (grey bars), pooling males and females together. The black bar represents the mean phenotypic distance between sympatric species, the *p-value* is based in the number of simulations where the phenotypic distance between species is higher than the mean value of inter-specific distance observed in sympatry.

### Signal of convergent iridescence among closely-related sympatric *Morpho* species

We used a permutation approach applied on the colorimetric variables to test whether the iridescent signal is more similar among species within sympatric populations than expected at random (Figure 3.B). The average phenotypic distance between two heterospecific pairs of individuals was significantly smaller in sympatry than the null distribution simulated by randomising sympatric and allopatric localities (*p-value* = 0.0005). Except for the two Venezuelan localities (VB and VT), the iridescent colouration in interspecific comparisons were more similar within all sympatric localities. When analysing the iridescent signals of males and females extracted from the same morpho-space separately (Figure S4), similar results were obtained: the mean phenotypic distance between sympatric males (Distmales = 53) is similar to the wing phenotypic distance between females (Distfemales = 54). However, the null distribution of the phenotypic variance of the males (Figure 4.A) is lower than the null distribution of the phenotypic variance of the females (Figure 4.B), likely due to a higher divergence of iridescence of females in allopatry.

Furthermore, the quantification of the phenotypic variance of iridescence showed that the trace of the covariance matrix of sympatric *M. helenor* (*Tsymp* = 4584) is lower than the trace of the covariance matrix of allopatric *M. helenor* (*Tallo* = 7194), suggesting lower phenotypic variation in *M. helenor* populations living in sympatry with *M. achilles*. Interestingly, the same trend is found for both sexes but the variance of iridescence is higher in females than in males (Males: *Tsymp* = 4239 and *Tallo* = 5885; Females: *Tsymp* = 4778 and *Tallo* = 8468).

Note that a signal a phenotypic convergence in sympatry is still found between *M. helenor* and *M. achilles* when generating the null phenotypic distance distribution only with the populations of the Amazonian Basin (Figure S5).

The results of the permutation test using the whole reflectance dataset also yielded similar results (Figure S6), although Venezuelan species appear even more divergent than the rest of the sympatric localities.

### Discrimination of iridescent wing signals by avian predators

The visual discrimination of the wing iridescence of species in sympatry was estimated using a UV-sensitive avian visual model: we measured the chromatic and achromatic distances between the reflectance spectra obtained at different angles of illumination between *M. helenor*/*M. achilles* individual pairs sampled in the same localities (see Figure S7 for chromatic distances and Figure S8 for achromatic distances). This discrimination dataset is summarised in Table 1. The least convergent *M. helenor*/*M*.*achilles* pairs distributed in the two Venezuelan localities (VT and VB) are highly discriminated by the modelled visual system of bird observers, especially at the achromatic level. In contrast, most of the significantly convergent populations were not discriminated at most angular positions. However, in some sympatric areas of Peru (PMH and PHH), Bolivia (LP) and Brazil (BCA), bird observers might be able to discriminate slight chromatic and achromatic differences at some specific angles of observations.

**Table 1:**
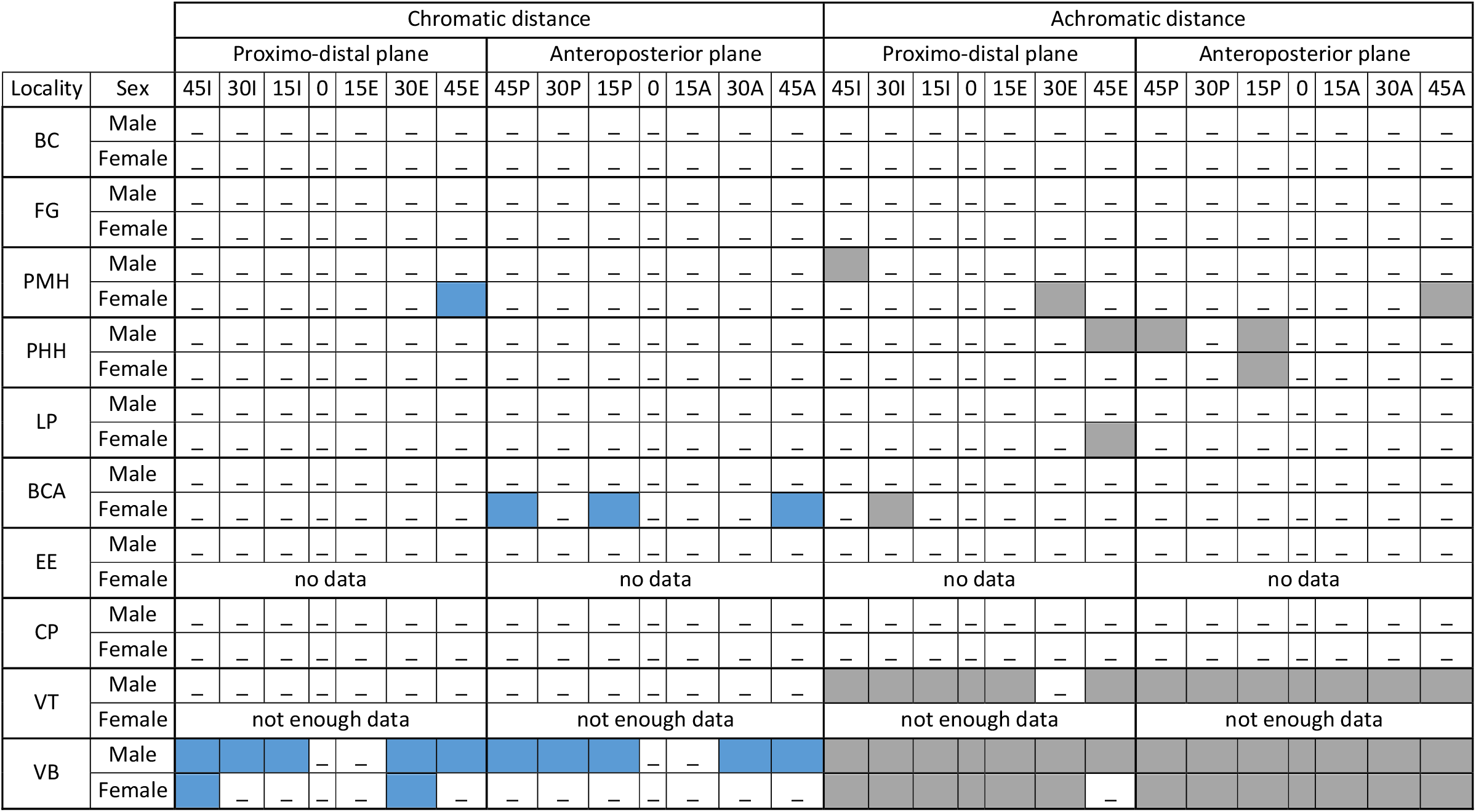
**Outline of the visual discrimination analyses of sympatric *Morpho* wings by a UV-sensitive bird** created from the results presented in Figure S5 and S6. The chromatic and achromatic distances of the wing reflectance of sympatric *Morpho* species were measured on the proximo-distal and antero-posterior plane of the wings at different angles of illumination, using a UV-sensitive bird visual model, for *M. helenor* and *M. achilles* males and females separately, within each sympatric locality. The coloured tiles highlight the groups for which the mean chromatic or achromatic distance (including its confidence interval) at a specific angle is higher than 1 *Just Noticeable Difference* (JND), used as a threshold of visual discrimination. Significant chromatic distances are marked in blue, while significant achromatic distances are marked in grey. The localities among which a signal of convergent iridescence has been detected are bolded

## Discussion

The comprehensive quantification of the structural colouration of 351 specimens of *Morpho* butterflies allowed to detect a significant signal of repeated convergent evolution of iridescence in multiple geographic areas where the sister species *M. helenor* and *M. achilles* occur in sympatry. The high similarity in the iridescence of the dorsal blue band in sympatry likely reinforces the visual resemblance between individuals from these two species, whose dorsal patterning was previously documented as locally convergent (Llaurens et al., 2021).

### Convergent iridescence in sympatry: the consequence of local abiotic factors?

The convergent evolution of traits in sympatric species is likely due to strong local selective pressures. What could be the main selective forces involved in iridescence convergence in sympatric *Morpho* sister species? First, shared environmental conditions and abiotic factors are expected to promote similar local adaptations between species living in the same habitat (MacLean et al., 2016; Siemers et al., 2024). In sympatry, *M. helenor* and *M. achilles* both live in similar micro-habitats in the forest understory, sharing identical light environments and temperatures within each locality, although flying at slightly different time periods (Bouinier et al., 2025; Le Roy et al., 2021). Thermodynamic tests have shown that iridescent butterfly scales may have a higher solar absorbance than pigmentary scales (Bosi et al., 2008), likely due to their light reflecting nanostructures, also involved in colour display. However, the precise link between the morphology of scale nanostructures and thermoregulation capacities is not clear: whether subtle nanostructure changes, triggering the reflection of different visual signals (Siddique et al., 2013) could also impact thermoregulation in iridescent blue *Morpho* is currently unknown. Furthermore, heat shock experiments showed that *Morpho* butterflies with iridescent scales have similar thermoregulation capacities compared to non-iridescent *Morpho* butterflies (Bouinier et al., 2025), suggesting that iridescent scales are not necessarily involved in temperature resistance in *Morpho*, and that other factors may impact the evolution of this trait.

### The repeated evolution of convergent iridescent signals is consistent with the evasive mimicry hypothesis

We repeatedly found a high resemblance in iridescence among individuals from *M. helenor* and *M. achilles* within several sympatric localities. Considering that current gene flow is currently stopped between these sympatric *Morpho* species (Le Roy et al., 2021), these iridescent phenotypic similarities are unlikely to result from repeated events of introgression, as observed in other mimetic butterfly species living in sympatry (Dasmahapatra et al., 2012). These similarities could also stem from shared ancestral polymorphism or independent evolution of similar phenotypic variations in these two species. In either case, the observed phenotypic convergence is consistent with a primary effect of evasive mimicry driven by the behaviours of the local communities of predators on colouration evolution. *Morpho* iridescence is a visual cue that could promote the escape of these butterflies against predators (Pinheiro, 1996; Vieira-Silva et al., 2024; Young, 1971) and facilitate predator learning via their conspicuousness (Pinheiro & Campos, 2019): these factors could have promoted the observed convergent evolution of iridescent signals via escape mimicry, especially of the hue variation of their wings. The reduced variance in iridescence within sympatric populations of *M. helenor* as compared to allopatric ones further suggests that phenotypic evolution within species might be constrained by phenotypic variation in other sympatric species. Interestingly, the variance of the iridescent signal was higher among females than among males. Because *Morpho* males typically patrol in open, exposed habitats where they are more visible to predators, whereas females usually remain concealed in the forest understory (Le Roy et al., 2021), males are likely subject to stronger predation pressures while other forces such as sexual selection might constrain the evolution of female wings. This may select for greater uniformity in the iridescent signal of males, leading to reduced phenotypic variance compared to females.

Although we found a repeated evolution of convergent iridescent colours in sympatry, the mean iridescent signal varied between localities. This geographic heterogeneity of iridescent signals may be driven by local variation in predator communities, as found in *Heliconius* butterflies: variations in local predator communities are indeed identified as driving the emergence of different mimicry rings between *H. melpomene* and *H. erato* for example (Mallet, 2010; Mallet and Gilbert, 1995). Models show that predator perception, preferences and behaviours may promote the selection of diverse mimetic patterns in prey from different localities maintained by positive frequency-dependent selection (Aubier & Sherratt, 2015), as shown in butterflies but also in frogs (Symula et al., 2001), bees and wasps (Chatelain et al., 2023), or ants (Wilson et al., 2012).

By estimating the visual discrimination of *Morpho* iridescence using a vision model corresponding to UV-sensitive birds, we also showed that the minute differences observed in sympatric areas between individuals from different species are unlikely to be detected by birds in most geographic areas. Iridescent wing colouration exhibited by the two sympatric species of *Morpho* are thus perceived as visually highly similar by birds, consistent with the hypothesis of evasive mimicry.

### Direct and indirect effect of iridescence on predator attacks

Differences between species in reflectance displayed at certain angles are nevertheless observed in some localities where convergence in iridescence was detected, highlighting that the local convergence can be somewhat limited. Such limitation could be due to divergent developmental constraints in the two species or differences in the available standing genetic variations causing iridescence evolution in the two species. Furthermore, even if detected by birds, minute differences in reflectance at certain angles might not prevent bird generalisation and avoidance, as shown by the prevalence of imperfect mimicry among aposematic prey (Kikuchi & Pfennig, 2013). For instance, predators tend to generalise aposematic signals and be less sensitive to slight phenotypic variation of aposematic prey in communities exhibiting more diverse visual signals (Ihalainen et al., 2012), allowing the evolution of non-accurate mimicry. The iridescent blue colour of *Morpho* being relatively rare within the highly diverse prey community they live in, the minute differences in iridescence between species, even if perceived by predators, may not be significant enough to prevent generalisation.

Furthermore, in *Morpho* butterflies, iridescence directly participates in predator escape due to the confusion effect limiting the catching success in case of predator attack. Selection for accurate mimicry between iridescent evasive species might be less stringent than for aposematic prey displaying pigmentary colours where colour itself has no effect on survival during the attack. Such direct survival advantage provided by iridescence during predator attack may thus limit the survival advantage brought by mimicry and thus weaken the selection on resemblance. Identifying what part of the iridescent signal generates confusion and is memorised by predators is now needed to assess how predator behaviour affects the evolution of iridescent wing pattern in *Morpho*. In *Morpho*, the relative importance of flashes during flight or colour variation as evasive/dissuasive signals for predators has not been studied. For instance, dynamic colours might reduce predation success when a prey is moving (Murali, 2018), as well as flashing signals involving rapid changes of brightness (Loeffler-Henry et al., 2021): what could be the incidence of these effects combined in *Morpho* butterflies? Whether all the colours displayed on the iridescent spectrum of *Morpho* butterflies are important in predator confusion and learning is still an open question.

Interestingly, no signal of phenotypic convergence was found between the sympatric *M. helenor* and *M. achilles* sister species in two localities in Venezuela (VT and VB). This result is consistent with those of Llaurens *et al*. 2021, showing that the width of their blue band also diverges in these populations. This result could be due to as possibly too large sampling delimitation, pooling together different populations submitted to different predator communities. Nevertheless, Venezuela is also a contact zone between populations usually isolated by the Andes (Blandin & Purser, 2013) and therefore may have retained larger phenotypic variability as a crossroad between divergent sub-species of *M. helenor*.

## Conclusion

Our results are consistent with a primary effect of selection exerted by local predator communities on the evolution of convergent iridescent colourations between closely-related *M. helenor* and *M. achilles* living in sympatry. Behavioural experiments investigating the effect of variation in iridescence pattern on predators are now needed to validate this hypothesis. Our results also highlight the importance of quantifying the visual effect of hue variations vs. brightness variations in order to identify the evolutionary drivers of structural colouration in butterflies.

## Acknowledgments

J.L. PhD was funded by an IBEES grant from Sorbonne Université. This study was funded by the European Union (ERC-2022-COG - OUTOFTHEBLUE - 101088089). Views and opinions expressed are however those of the authors only and do not necessarily reflect those of the European Union or the European Research Council. Neither the European Union nor the granting authority can be held responsible for them.

## Data availability

The data and scripts used in this article will be available in a Zenodo repository upon publication

