## Supplementary files for "Repeated evolution of convergent iridescence in closely-related species of *Morpho* butterflies living in sympatry"

### Supplementary Information

Table S1: Number of specimens used for the wing reflectance measurements from the collection of the National Museum of Natural History (Paris, France). The subspecies name among each geographical locality is given according Blandin (2007), sympatric localities are highlighted in black and allopatric localities are highlighted in red.

| Biogeo-graphic region | Species co-occurrence | Sampling zone | Code | *M. helenor* | | | *M. achilles* | | | |
|---|---|---|---|---|---|---|---|---|---|---|
| | | | | Subspecies | ♂ | ♀ | Subspecies | ♂ | ♀ | Total |
| *trans-andean* | Allopatry | Central America | CA | *montezuma* | 6 | 6 | - | - | - | 12 |
| | Allopatry | Costa Rica | CR | *carillensis* | 6 | 6 | | | | 12 |
| | Allopatry | Costa Rica/Panama | CRP | *limpida* | 6 | 1 | - | - | - | 7 |
| | Allopatry | Western Colombia | WP | *microphthalmus* | 6 | 6 | - | - | - | 12 |
| | Allopatry | Central Colombia | CC | *peleides* | 6 | 1 | - | - | - | 7 |
| *cis*-Andean | Sympatry | Eastern Ecuador | EE | *theodorus* | 10 | 10 | *phokylides* | 6 | 0 | 26 |
| | Allopatry | Western Ecuador | WE | *bristowi* | 10 | 10 | - | - | - | 20 |
| | Allopatry | Peru Middle Marañon | PM | *charapensis* | 6 | 6 | - | - | - | 12 |
| | Sympatry | Peru Middle Huallaga | PMH | *theodorus* | 6 | 3 | *phokylides* | 6 | 2 | 17 |
| | Sympatry | Peru High Huallaga | PHH | *lacommei* | 6 | 5 | *fagardi* | 6 | 4 | 21 |
| | Sympatry | Central Peru | CP | *papirius* | 6 | 6 | *agamedes* | 6 | 6 | 24 |
| | Sympatry | Bolivia La Paz | LP | *coelestis* | 6 | 4 | *songo* | 6 | 6 | 22 |
| | Sympatry | Bolivia Cochabamba | BC | *coelestis* | 6 | 6 | *vitrea* | 6 | 6 | 24 |
| | Sympatry | Brazil Central Amazon | BCA | *helenor* | 6 | 6 | *achilles* | 6 | 1 | 19 |
| | Sympatry | Venezuela Bolívar | VB | *tepuina* | 6 | 5 | *glaisi* | 6 | 2 | 19 |
| | Sympatry | Venezuela Tucupita | VT | *tucupita* | 5 | 1 | *guaraunos* | 6 | 1 | 13 |
| | Allopatry | Venezuela El Pao | EP | *ululina* | 6 | 3 | - | | - | 9 |
| | Sympatry | Surinam/French Guiana/Northern Pará | SFR | *helenor* | 10 | 10 | *achilles* | 10 | 10 | 40 |
| Atlantic | Allopatry | Argentina Misiones | AM | *achillides* | 6 | 5 | - | - | - | 11 |
| | Allopatry | Brazil Rio de Janeiro | BR | *achillaena* | 6 | 6 | - | - | - | 12 |
| | Allopatry | Brazil João Pessoa | BJ | *anakreon* | 6 | 6 | - | - | - | 12 |
| Total | | | | | 137 | 112 | | 64 | 38 | **351** |

**A.**

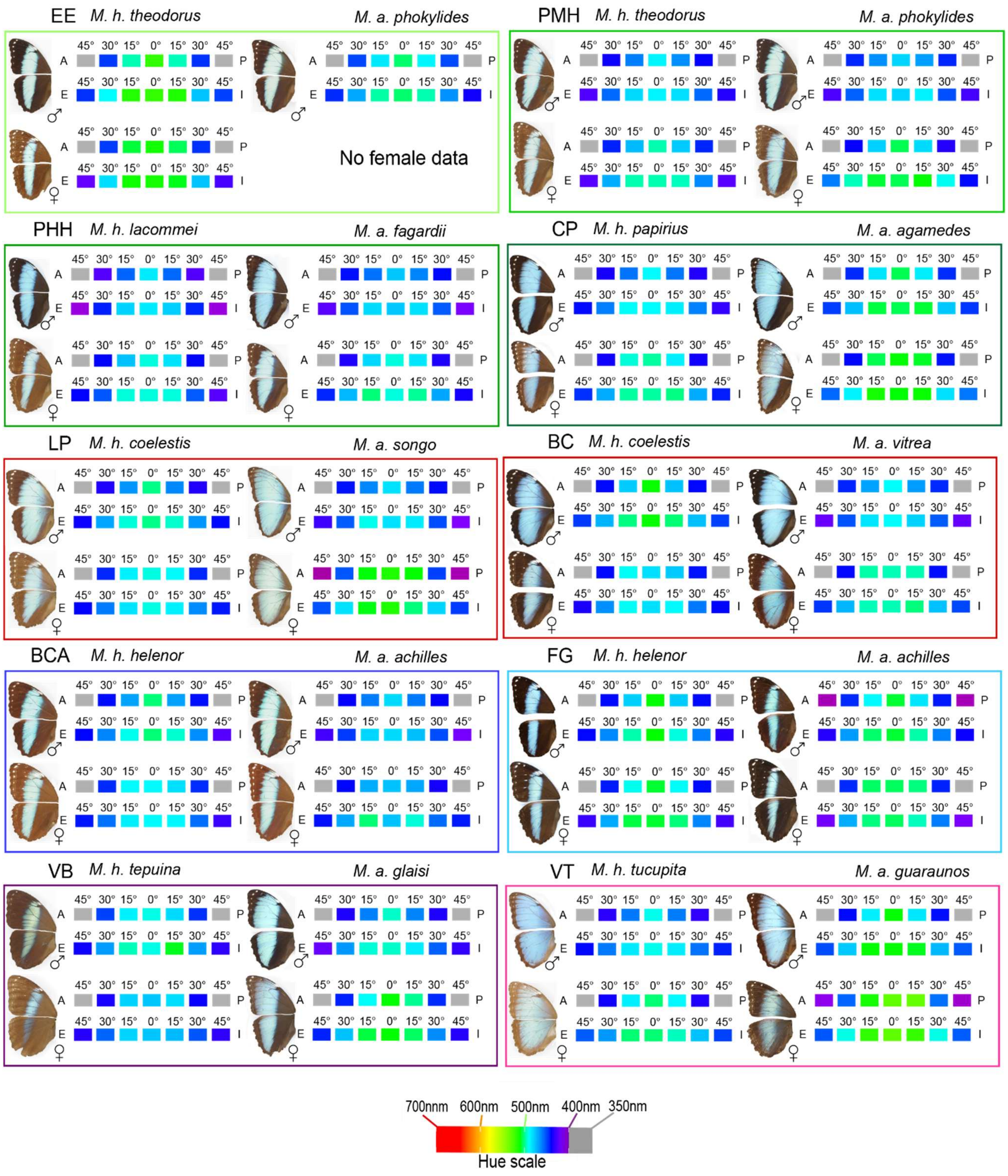
EE
M. h. theodorus
M. a. phokylides
No female data
PMH
M. h. theodorus
M. a. phokylides
PHH
M. h. lacommei
M. a. fagardii
CP
M. h. papirius
M. a. agamedes
LP
M. h. coelestis
M. a. songo
BC
M. h. coelestis
M. a. vitrea
BCA
M. h. helenor
M. a. achilles
FG
M. h. helenor
M. a. achilles
VB
M. h. tepuina
M. a. glaisi
VT
M. h. tucupita
M. a. guaraunos
45° 30° 15° 0° 15° 30° 45°
A
P
E
I
♂
♀
700nm 600nm 500nm 400nm 350nm
Hue scale

**B.**

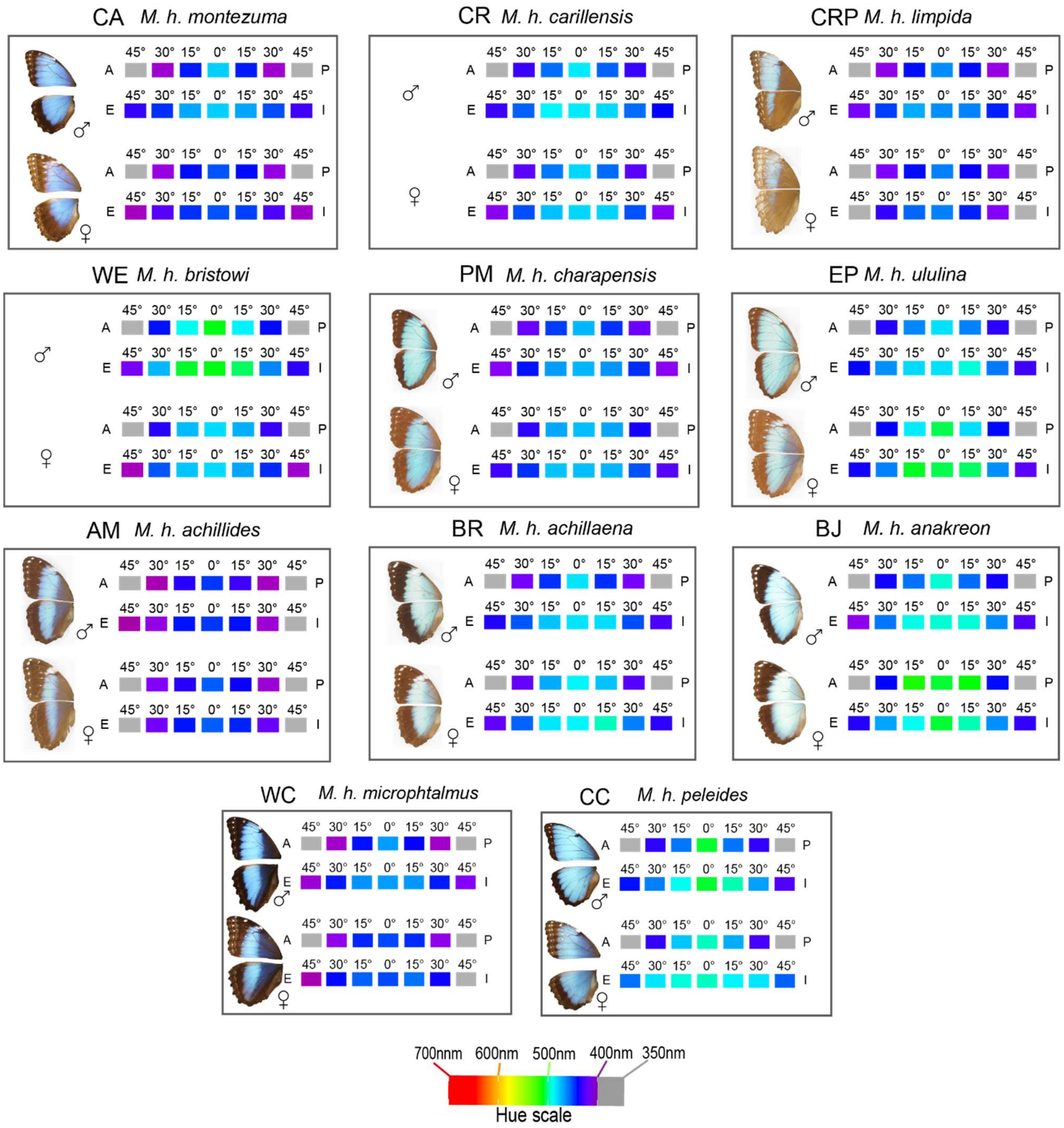


Figure S1: Hue iridescence measured among *M. helenor*, *M. achilles* and *M. granadensis* in (A) sympatric and (B) allopatric populations. Each box represents a studied locality, and contains the picture of male (top) and female (bottow) wing morphology belonging to sympatric sister species (among sympatric locations) or to *M. helenor* only (among allopatric locations). For each sex, the mean Hue value among individuals was calculated using the reflectance spectra measured at every measured studied angle, either on the Antero-posterior (first row) or on the Proximo-distal (second row) plane of the wings. Mean hues ranging from 400nm to 700nm are represented using a RGB color scale ranging from purple to red following the color variations observed on the visible light spectrum. Mean hues ranging from 350nm to 400nm in the UV-range are colored in grey. Note that this color representation is only illustrative and do not simulate butterflies’ nor predators’ vision.

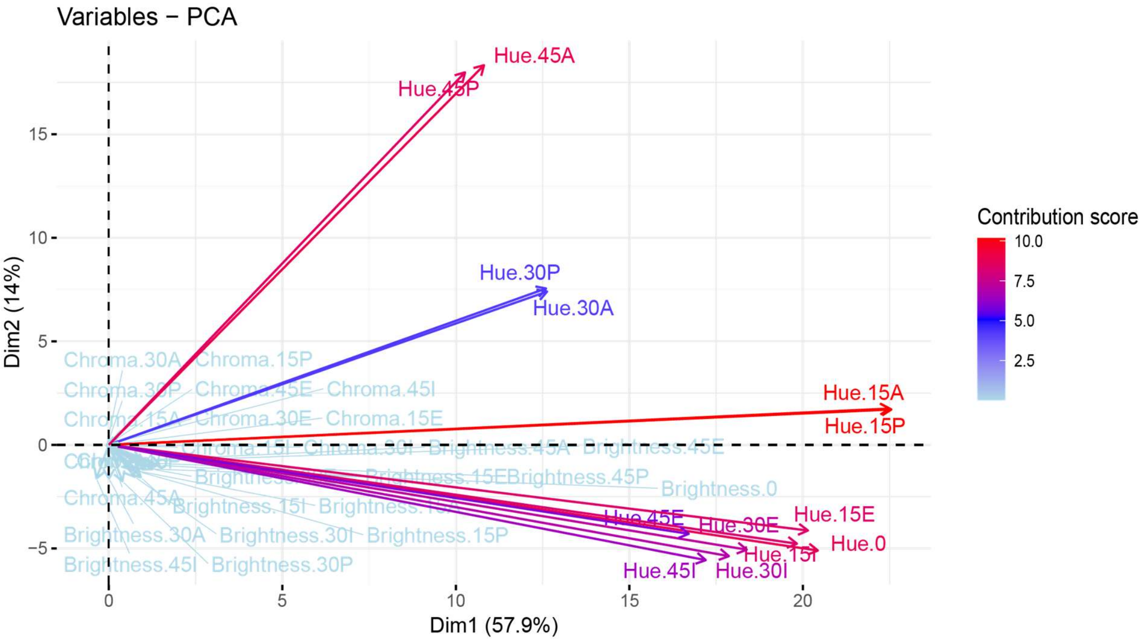


Figure S2: Visual representation of the contribution to the PCA representation of Hue, Brightness and Chroma variables measured on the Proximo-distal and Antero-posterior plane at different angles of illumination: both axes are correlated to Hue variables at different angles of illumination.

**A.**

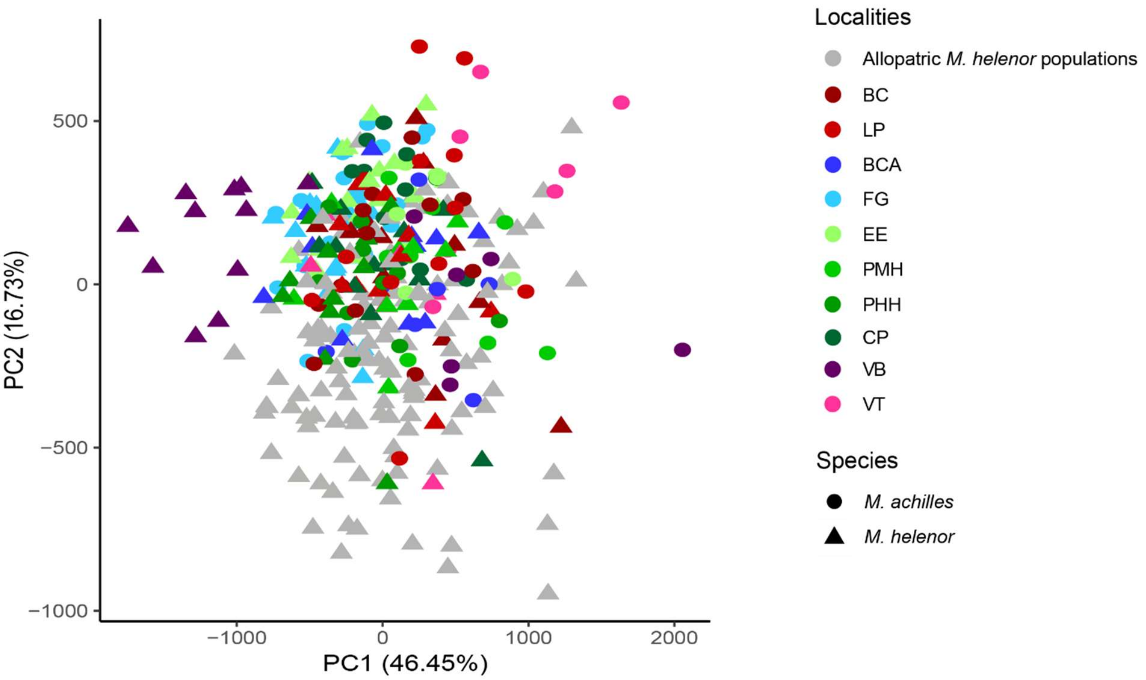


**B.**

| Variables | Df | SumOfSqs | R2 | F | Pr(>F) |
|---|---|---|---|---|---|
| Species | 1 | 7121217 | 0.04256886 | 27.138524 | **0.001**** |
| Sex | 1 | 9215814 | 0.05508984 | 35.120908 | **0.001**** |
| Locality | 20 | 45799237 | 0.27377643 | 8.726906 | **0.001**** |
| Species*Sex | 1 | 1782905 | 0.01065776 | 6.794544 | **0.002**** |
| Species*Locality | 9 | 13959759 | 0.08344796 | 5.911088 | **0.001**** |
| Sex*Locality | 20 | 10905046 | 0.06518765 | 2.077923 | **0.001**** |
| Species*Sex*Locality | 8 | 2406292 | 0.01438421 | 1.146279 | 0.281 |
| Residual | 290 | 76096725 | 0.45488727 | NA | NA |
| Total | 350 | 167286996 | 1 | NA | NA |

Figure S3: (A) Principal component analysis based on the "whole reflectance" dataset, using the reflectance of every nanometer for each angle of illumination as a variable, accounting for the variation of colouration of *Morpho* butterflies on both wing planes. Each point on this graph represents the global iridescence signal of 1 individual, while considering angle variation: the closer the points are together, the more similar the iridescence of two individuals is. Grey points represent the iridescent signal of allopatric *M. helenor* found living in allopatry, while *Morpho* individuals sharing their habitat with closely-related species are highlighted in colors according to their locality shown on Figure 1. *M. achilles* individuals are represented with circles, and *M. helenor* with triangles. Note that PC1 and PC2 account for more than 60% of the variance. (B) PERMANOVA results testing for the effect of Species, Sex, Locality and the interactions between those variables on the first 10 PCs of the "whole reflectance" PCA. Similar to the PERMANOVA performed on the colorimetric variable coordinates in the main text, this method allows to detect a significant effect of species, sex, locality, and of the interactions between those terms, suggesting that iridescence is highly diversified, both geographically and between species and subspecies.

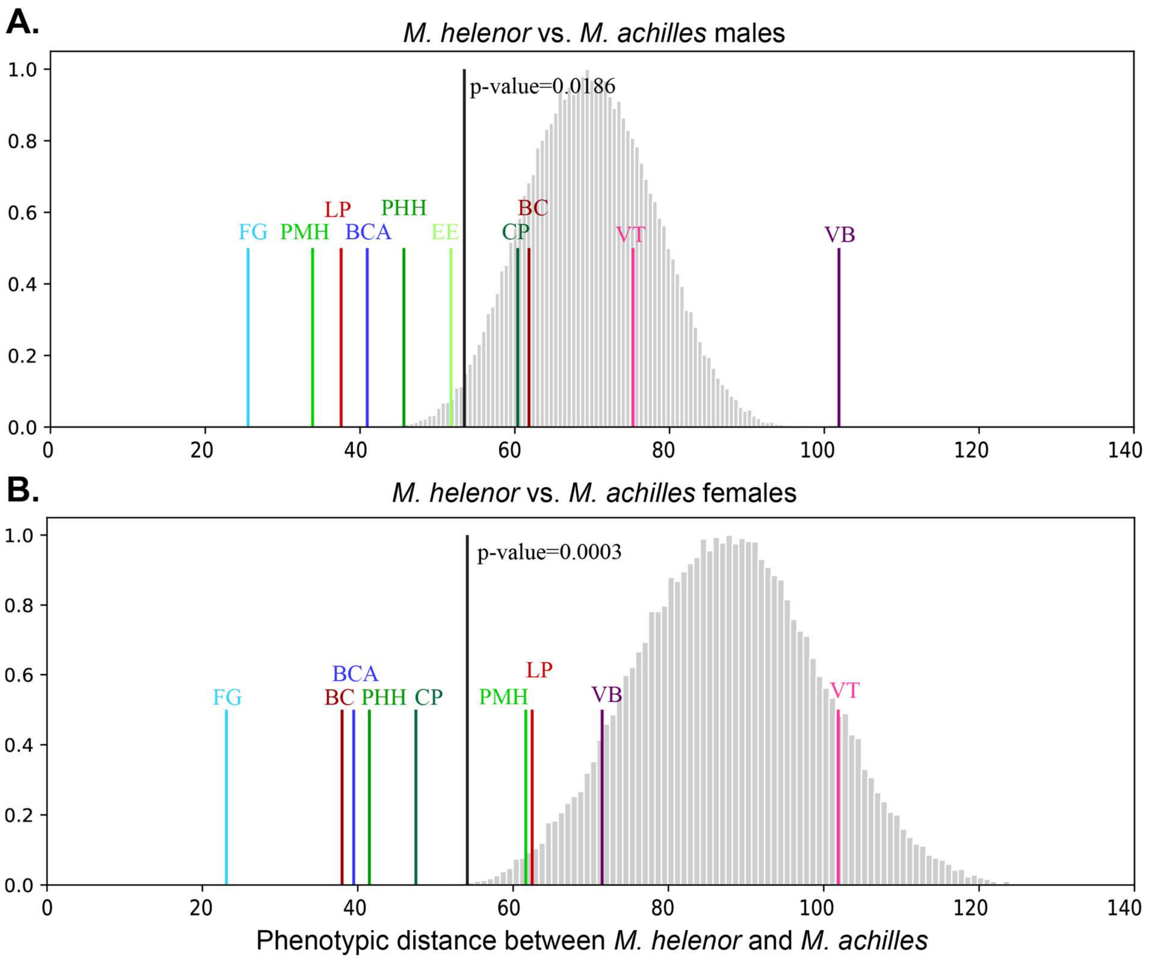


Figure S4: Phenotypic distances measured using the coordinates based on the Brightness-Hue-Chroma dataset, between *M. helenor* and *M. achilles* sharing the same locality (coloured bars) and the predicted null phenotypic distribution between species obtained using 10,000 bootstraps, randomly reallocating the different sampling zone between sympatric and allopatric species (grey bars). Graph (B) presents the global phenotypic distances measured by pooling males together, while graph (B) presents the global phenotypic distances measured by pooling females together. The black bar represents the mean phenotypic distance between sympatric species, the p-value is based in the number of simulations where the phenotypic distance between species is higher than the mean value of inter-specific distance observed in sympatry.

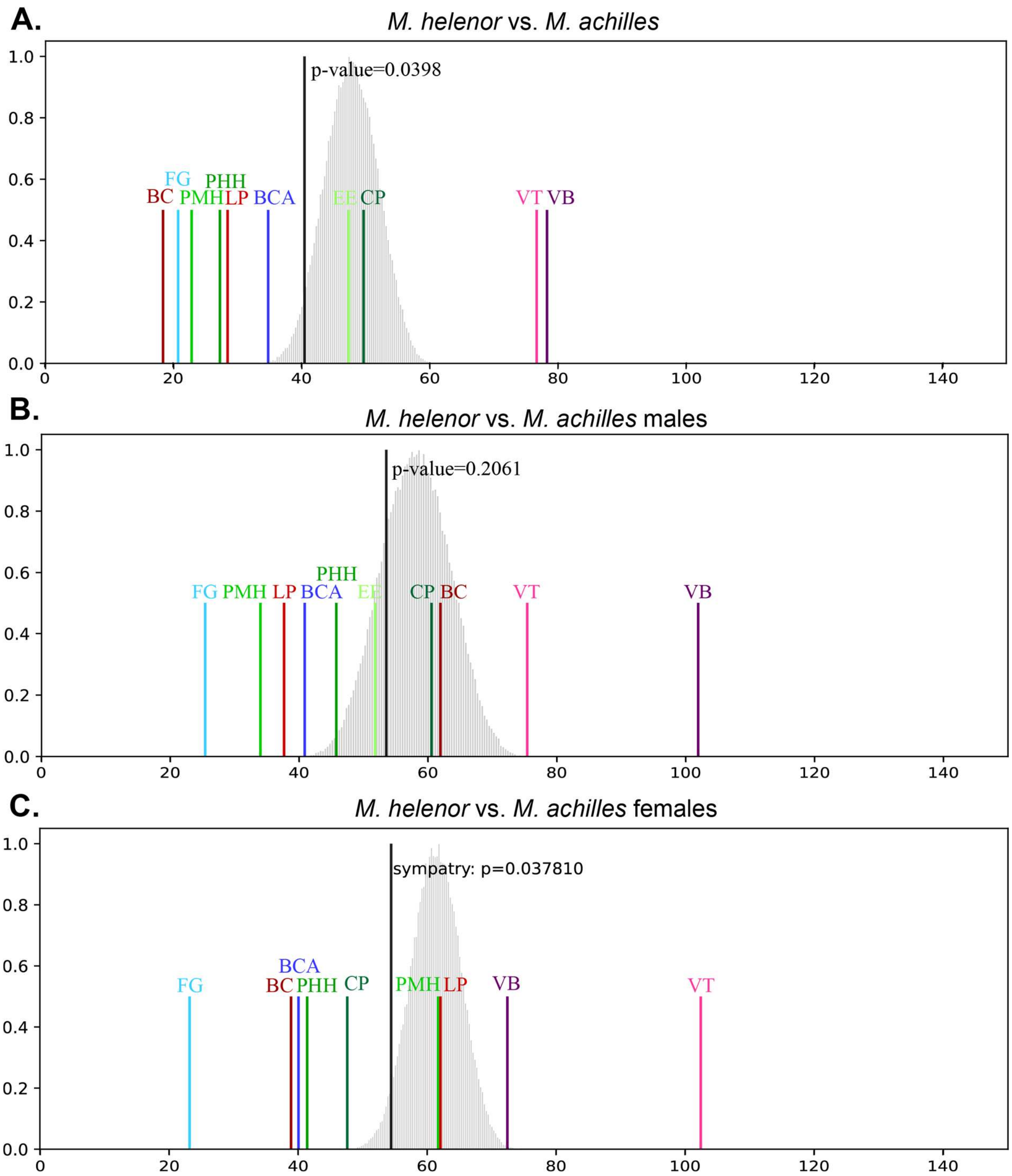


Figure S5: Phenotypic distances between M. helenor and M. achilles measured using the Brightness-Hue-Chroma dataset but only keeping the populations found in the Amazonian Basin (10 sympatric locations: FG, PMH, LP, BCA, PHH, EE, CP, BC, VT, VB; 2 allopatric locations where M. helenor is found isolated: PM and EP). The predicted null phenotypic distribution between species obtained using 10,000 bootstraps, randomly reallocating the different sampling zone between sympatric and allopatric species are shown in grey, and the phenotypic distance of each sympatric locations are shown in colours. Graph (A) present the global phenotypic distances measured by pooling males and females together, while graphs (B) and (C) only display the phenotypic distances measured between males and females only respectively. The black bar represents the mean phenotypic distance between sympatric species, the p-value is based in the number of simulations where the phenotypic distance between species is higher than the mean value of inter-specific distance observed in sympatry.

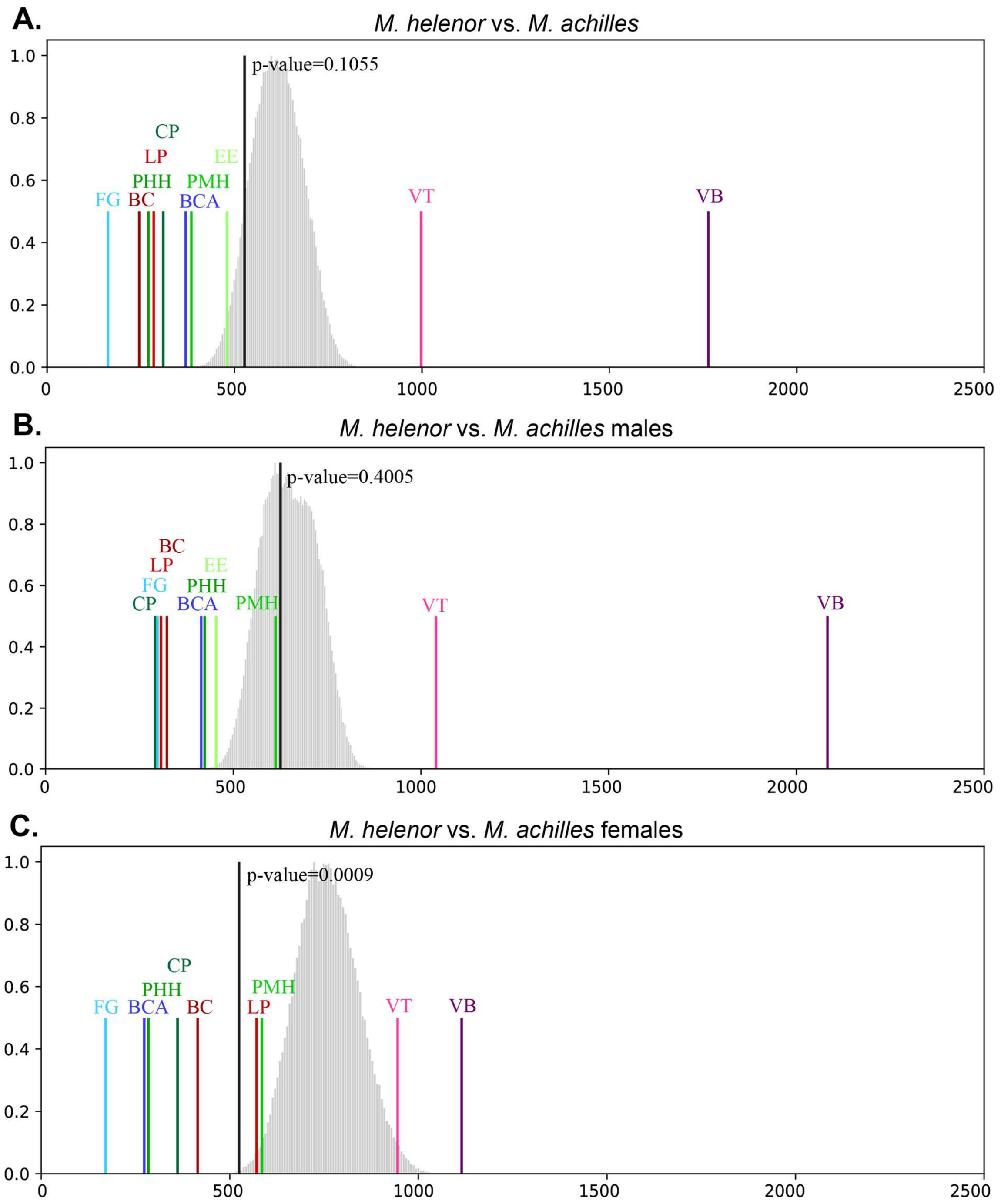


Figure S6: Phenotypic distances measured using the "whole reflectance" coordinates, between *M. helenor* and *M. achilles* sharing the same locality (coloured bars) and the predicted null phenotypic distribution between species obtained using 10,000 bootstraps, randomly reallocating the different sampling zone between sympatric and allopatric species (grey bars). Graph (A) present the global phenotypic distances measured by pooling males and females together, while graphs (B) and (C) only display the phenotypic distances measured between males and females only respectively. The black bar represents the mean phenotypic distance between sympatric species, the p-value is based in the number of simulations where the phenotypic distance between species is higher than the mean value of inter-specific distance observed in sympatry. Compared to the permutation results obtained with the colorimetric variables dataset, this method detects a greater divergence of *M. helenor* and *M. achilles* iridescent colour patterns in Venezuela, strongly influencing the mean phenotypic distance between sympatric species (black bar). However, a convergence of iridescent signal is still detected in the other sympatric populations. Furthermore, phenotypic resemblance is higher in sympatric females than in sympatric males.

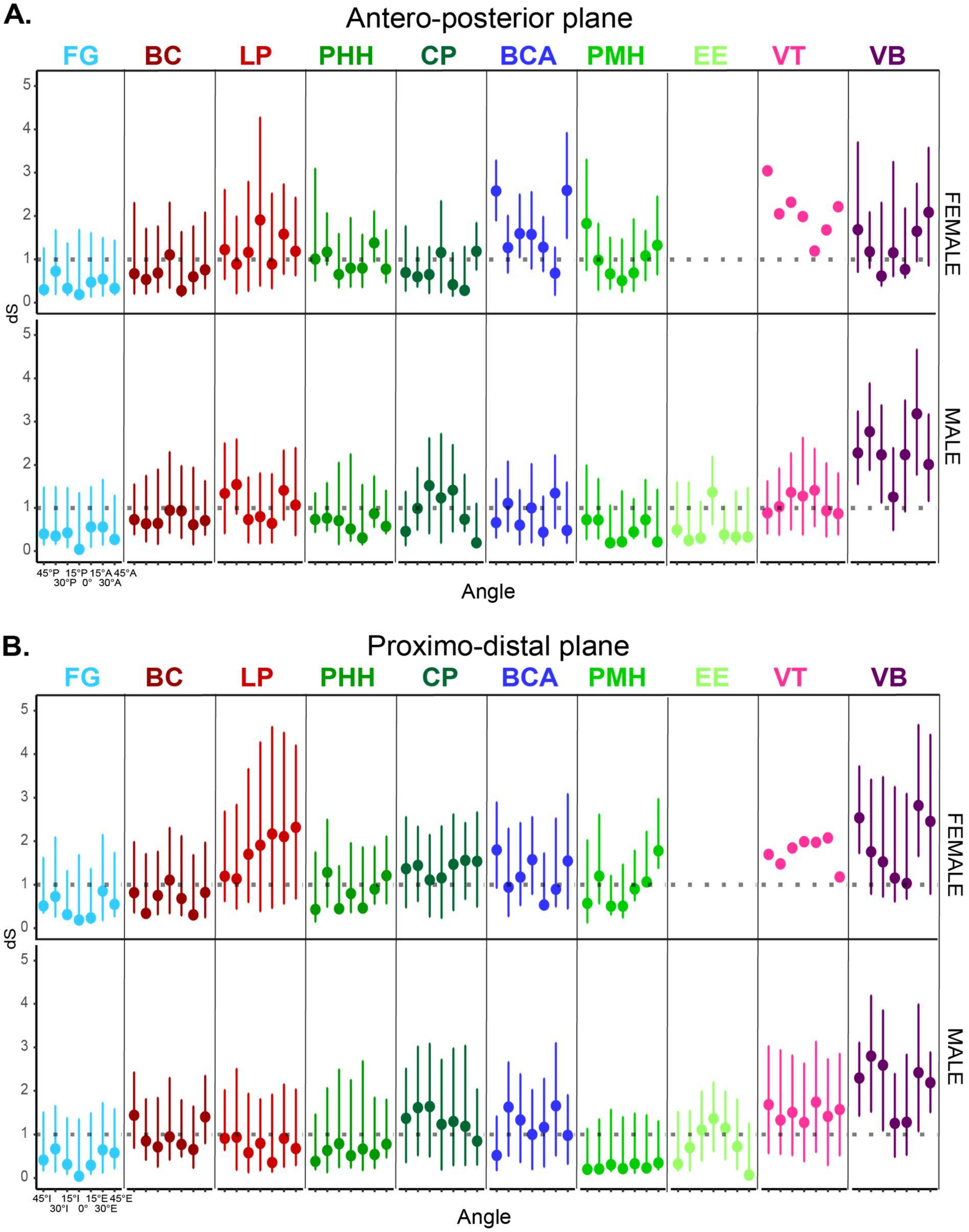


Figure S7: Chromatic distances (i.e. the visual discrimination rate of hue by an avian visual model) of the wing reflectance measured (A) on the Antero-posterior and (B) the Proximo-distal planes of Morpho wings. The chromatic distances have been measured separately between females (first row of each figure) and males (second row of each figure) of the two species sharing each sympatric location, in order to account for sexual dimorphism. Among each locality, the tested angles are displayed from right to left following the order: [-45°, -30°, -15°, 0°, 15°, 30°, 45°] on the Antero-posterior plane, and [-45°I, -30°I, -15°I, 0°, 15°, 30°, 45°] on the Proximo-distal plane. The dotted line represents the threshold of discrimination of JND=1: visual discrimination is considered possible if the measured the mean chromatic distance (coloured circle) and its confidence interval (coloured vertical line) are superior to the threshold.

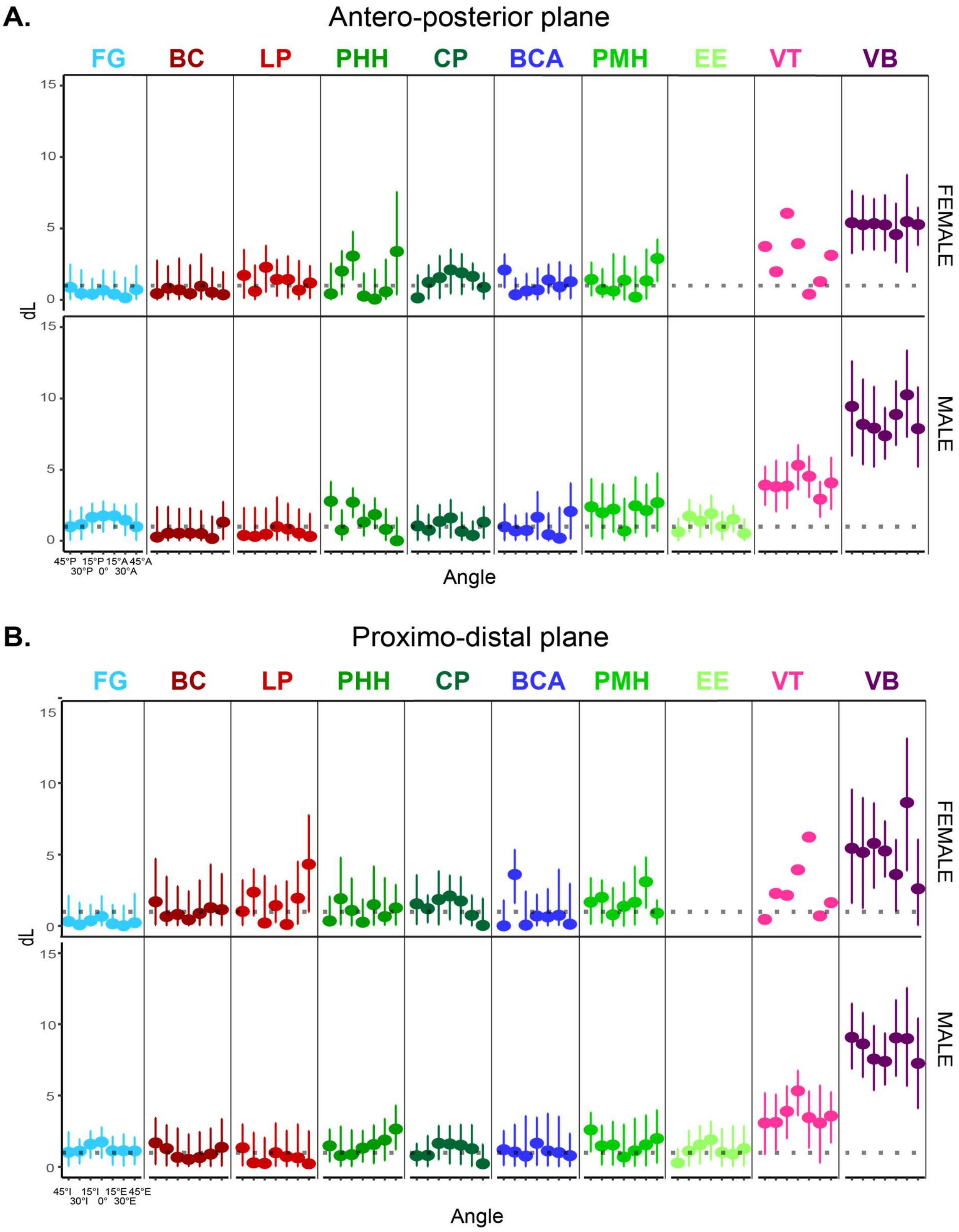


Figure S8: Achromatic distances (i.e. the visual discrimination rate of brightness by an avian visual model) of the wing reflectance measured (A) on the Antero-posterior and (B) the Proximo-distal planes of Morpho wings. The achromatic distances have been measured separately between females (first row of each figure) and males (second row of each figure) of the two species sharing each sympatric location, in order to account for sexual dimorphism. Among each locality, the tested angles are displayed from right to left following the order: [-45°, -30°, -15°, 0°, 15°, 30°, 45°] on the Antero-posterior plane, and [-45°, -30°, -15°, 0°, 15°, 30°, 45°] on the Proximo-distal plane. The dotted line represents the threshold of discrimination of JND=1: visual discrimination is considered possible if the measured the mean achromatic distance (coloured circle) and its confidence interval (coloured vertical line) are superior to the threshold.
